# Multi-omics definition of the sex-specific glycoproteome of murine tissues

**DOI:** 10.64898/2026.03.10.710926

**Authors:** Rebeca Kawahara, Masaya Hane, Di Wu, Bingyuan Zhang, Fumiya Sakamoto, Takahiro Nakagawa, Takayuki Omoto, Kristina Mae Bienes, Naaz Bansal, Zeynep Sumer-Bayraktar, Sayantani Chatterjee, Koichi Himori, Chiaki Nagai-Okatani, Atsushi Kuno, Makoto Kashima, Daniel Kolarich, Yusuke Matsui, Ken Kitajima, Kenji Kadomatsu, Chihiro Sato, Morten Thaysen-Andersen

## Abstract

Sex-specific differences in the glycoproteome remain poorly defined despite growing evidence that protein glycosylation is a key determinant of sex biology. Here we present a tissue-resolved glycoproteome atlas of adult male and female C57BL/6J mice, integrating transcriptomics, proteomics and glycoproteomics with sialic acid speciation and lectin microarray profiling across 19 tissues. Quantitative analysis of >26,800 protein- and site-specific N-glycoforms from 1,512 glycoproteins revealed highly distinct tissue glycoproteomes shaped by coordinated regulation of protein abundance and glyco-enzyme expression. Multi-omics integration identified strong glycophenotype-enzyme relationships, including control of tissue sialylation by Cmas and Cmah, suggesting rate-limiting roles in glycosylation. Pronounced sex-linked glycophenotypes were observed in salivary gland, liver and kidney, driven by differences in fucosylation, sialylation and protein abundance, whereas the brain glycome was largely conserved between sexes. An interactive online database (https://igcore.cloud/mta/atlas-viewer/) provides a resource for exploring sex-biased glycosylation across mouse tissues.

## Introduction

Biological sex is a fundamental determinant of human physiology and disease, influencing immune function^1^, diseases susceptibility^2^, drug response and clinical outcomes^3^ across a wide spectrum of conditions. While many sex differences have been attributed to hormonal regulation and spatiotemporal expression of sex chromosome–encoded genes^4^, post-translational modifications remain comparatively underexplored in sex biology despite their well-recognized impact on protein function^5^. Among these, protein asparagine (*N*)-linked glycosylation, an abundant and structurally diverse type of post-translational modification, has emerged as a central regulator of protein stability, trafficking, signaling, and cell–cell communication^6,7^.

The *N*-glycoproteome is generated by a complex biosynthetic machinery integrating enzymatic processing, substrate availability, and secretory pathway organization. Following *en bloc* glycan transfer in the endoplasmic reticulum (ER), nascent *N*-glycoproteins undergo sequential trimming and remodeling during protein maturation, with further diversification in the Golgi through the concerted action of glycosidases and glycosyltransferases to produce molecular diversity in the form of glycan types (oligomannose, hybrid, complex, paucimannose) and terminal features such as antennary branching, sialylation and fucosylation^8,9^. This structural glycan diversity underpins tissue specialization, as glycosylation traits such as linkage-specific sialylation and fucosylation patterns are known to regulate molecular recognition, immune interactions, and protein function in an organ-dependent manner. Despite still poorly understood, emerging literature suggest that the tissue-specific expression, localization, and enzymatic activity of the glycosylation machinery are key contributors to the distinct, site-specific glycan signatures observed across tissues^10–12^. Enabled by advances in mass spectrometry–based glycomics and glycoproteomics^13^, comprehensive maps of *N*-glycans, providing both detailed structural resolution and site-specific information on protein carriers, have been generated across tissues^14,15^ and disease contexts^16–18^. Notably, recent reports have also demonstrated the impact of biological sex in shaping the plasma *N*-glycoproteome of human and mouse^19–22^ and in other organisms^23,24^. While these studies have greatly expanded our understanding of the glycosylation landscapes, the extent to which the heterogenous glycoproteome is shaped by biological sex within distinct tissue environments, and the regulatory mechanisms underlying such sex differences, remains unknown.

In this study, we present a comprehensive, multi-omics characterization of the *N*-glycoproteome in adult male and female C57BL/6J mice across tissues, providing new insight into how biological sex shapes tissue glycosylation. By integrating (glyco)proteomics and transcriptomics with sialic acid speciation and lectin microarray analyses, we demonstrate that the presentation of glycan structures on individual protein carriers is orchestrated by a complex, tissue-specific network of sialylation and fucosylation enzymes, in concert with a sex-biased proteome repertoire, particularly in the salivary gland, liver, and kidney. Lectin array-based profiling combined with porous graphitized carbon (PGC) LC-MS/MS glycomics further uncovered the linkage-specific fucosylation and sialylation modifications that define the glycophenotypes of these three sex-biased tissues. Notably, this study also reveals that the brain exhibits extensive glycome diversity that is largely conserved between male and female mice, highlighting constraints on sex-linked variations in glycosylation. To facilitate discovery, we have established an interactive database (https://igcore.cloud/mta/atlas-viewer/) that enables exploration of protein- and site-specific glycosylation across male and female murine tissues, providing a resource for investigating sex-specific glycoproteome variations and links to health and disease. Collectively, this work positions glycosylation as a critical determinant of tissue physiology, with implications for understanding mechanisms driving sex disparities in disease susceptibility, outcomes and response to treatment.

## Results

### Integrated multi-omics approach to define sex-linked glycophenotypes across mouse tissues

In this study, we applied a multi-omics approach to generate a sex-stratified *N*-glycoproteome atlas across 17 non-sex tissues and two sex tissues (testis [here shortened to TE, tissue #17] and ovary [OV, #18]) from adult male and female C57BL/6J mice (**Figure 1a**). To profile complementary molecular layers underlying the sex-specific glycoproteome diversity across murine tissues, the multi-omics workflow included comparative proteomics, glycoproteomics and transcriptomics integrated with three types of glycomics (PGC-LC-MS/MS- and lectin microarray [LMA]-based glycome profiling and HPLC-based sialic acid speciation analysis) (**Figure 1b**).

**Figure 1.**
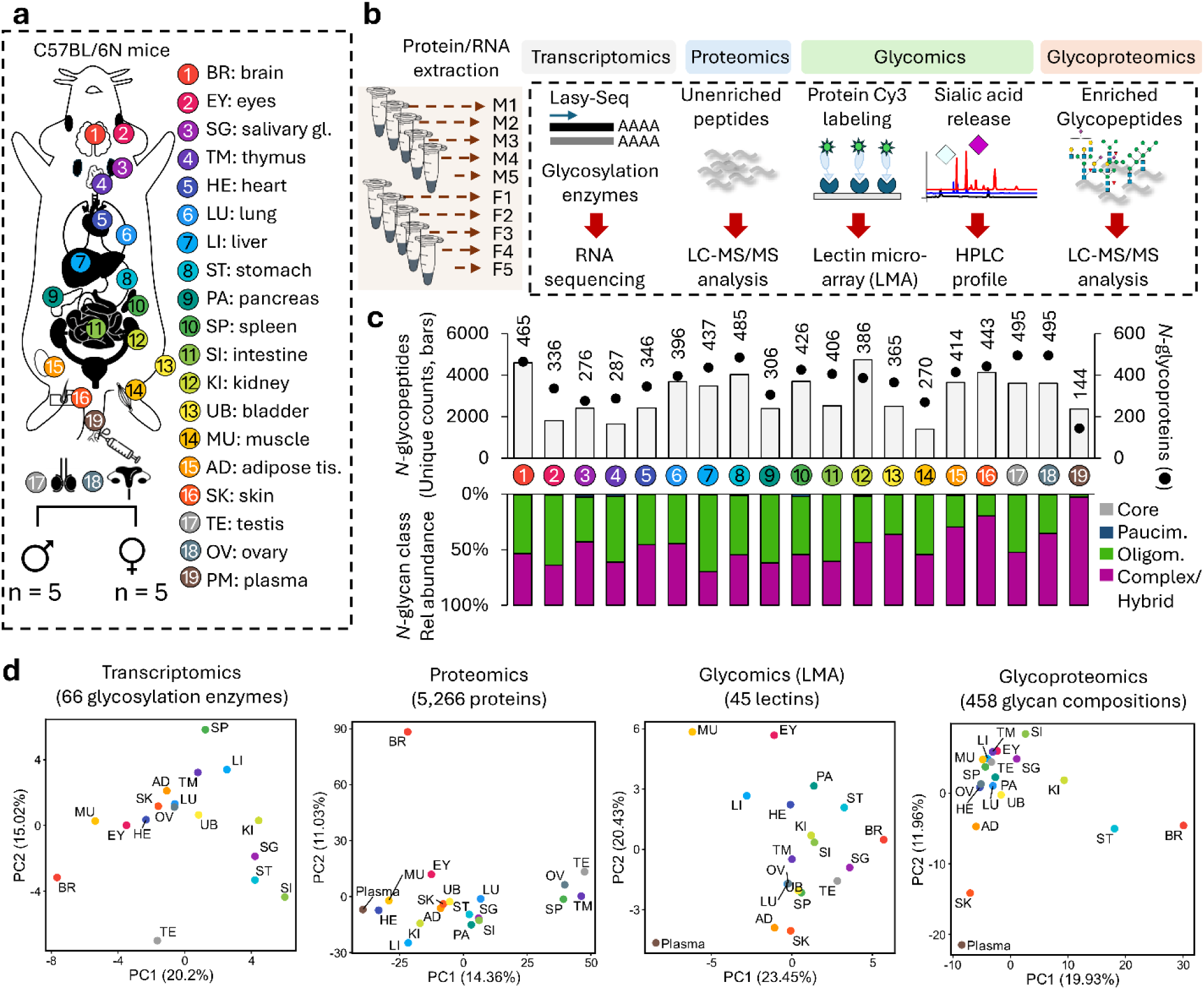
A sex-stratified mouse tissue *N*-glycoproteome atlas. **a**) The mouse *N*-glycoproteome was profiled across 17 non-sex-specific and 2 sex-specific (testis and ovary) tissues from adult C57BL/6J mice (10 weeks old, n = 5 females and n = 5 males). **b**) Overview of the applied multi-omics workflows spanning quantitative glycoproteomics, proteomics, and transcriptomics integrated with three types of glycomics including lectin microarray (LMA) and sialic acid speciation analyses as well as (for a select set of tissues) PGC-LC-MS/MS. **c**) Unique *N*-glycopeptides (bars), source *N*-glycoproteins (dots), and relative distribution of *N*-glycan types (mirror bars) identified by LC-MS/MS glycoproteomics across tissues, see key. **d**) Principal component analysis (PCA) showing tissue-specific molecular signatures across the multi-omics datasets.

Using a TMT-based (label-assisted) quantitative glycoproteomics approach, more than 26,800 unique protein- and site-specific *N*-glycoforms spanning ∼1,500 unique *N*-glycoproteins were identified and quantified, providing a comprehensive characterization of the male and female glycoproteomes of the 19 adult mouse tissues (**Supplementary Table S1**).

Each tissue exhibited a distinct *N*-glycoproteome shaped by differences in glycoprotein repertoires and variations in the distribution and relative abundance of site-specific glycoforms, predominantly composed of complex- and oligomannosidic-type *N*-glycans (**Figure 1c**). Despite considerable variations, more than 2,000 unique glycopeptides mapping to ∼250 different proteins were detected for most tissues with a particularly high glycoproteome coverage achieved for the brain (BR, tissue #1, 4,599 glycopeptides across 1,001 glyco-sites from 465 glycoproteins). In terms of molecular diversity, the brain (371 glycan compositions), stomach (ST, #8, 371 compositions), and kidney (KI, #12, 346 compositions) displayed the greatest glycome diversity whereas muscle (MU, #14, 158 compositions), thymus (TM, #4, 185 compositions), and eye (EY, #2, 190 compositions) showed the lowest glycome diversity. The tissue diversity in glycosylation was recapitulated when using multiple measures such as micro- and macro-heterogeneity and occupied glyco-sites per protein to determine the relative glycoproteome heterogeneity of each tissue (**Supplementary Figure S1a**). The kidney and brain exhibited the highest glycoproteome diversity while eye and muscle were less heterogenous tissues in term of *N*-glycosylation.

Multivariate biplots using the entire glycoproteomics data as input showed that oligomannose and terminal glycosylation features on complex *N*-glycans such as sialylation are principal glycoforms that contribute to tissue variations (**Supplementary Figure S1b**). These trends were strengthened by the LMA data showing that lectins against oligomannose (e.g. ConA, NPA), sialylation (e.g. SNA, SSA) and fucosylation (AAL) can separate mouse tissues.

Moreover, the mRNA expression profiles of 66 glyco-enzymes (glycosyltransferases and glycoside hydrolases) as measured by transcriptomics (**Supplementary Table S2**), the 5,266 proteins measured by proteomics (**Supplementary Table S3**), and the high level glycome data collected using 45 lectins recognizing diverse glycan epitopes, **Supplementary Table S4**) collectively reinforced tissue-specific molecular characteristics, with plasma and the brain consistently featuring distinctly different glycoproteomes across all layers (**Figure 1d**).

### Tissue-specific glycosignatures across cellular compartments

When considered at the global level (combined across all tissues), the mouse *N*-glycoproteome, as expected, was found to be highly enriched in glycoproteins annotated to localize to the extracellular space (adj. *p* = 5.16 x 10^-96^) and plasma membrane (adj. *p* = 3.80 x 10^-37^), with additional representation of proteins annotated to reside in the endoplasmic reticulum (ER, adj. *p* = 1.15 x 10^-36^), lysosome (adj. *p* = 2.37 x10^-22^) and Golgi apparatus (adj. *p* = 1.02 x 10^-04^) and other vesicular compartments (**Figure 2a** and **Supplementary Table S1**).

**Figure 2.**
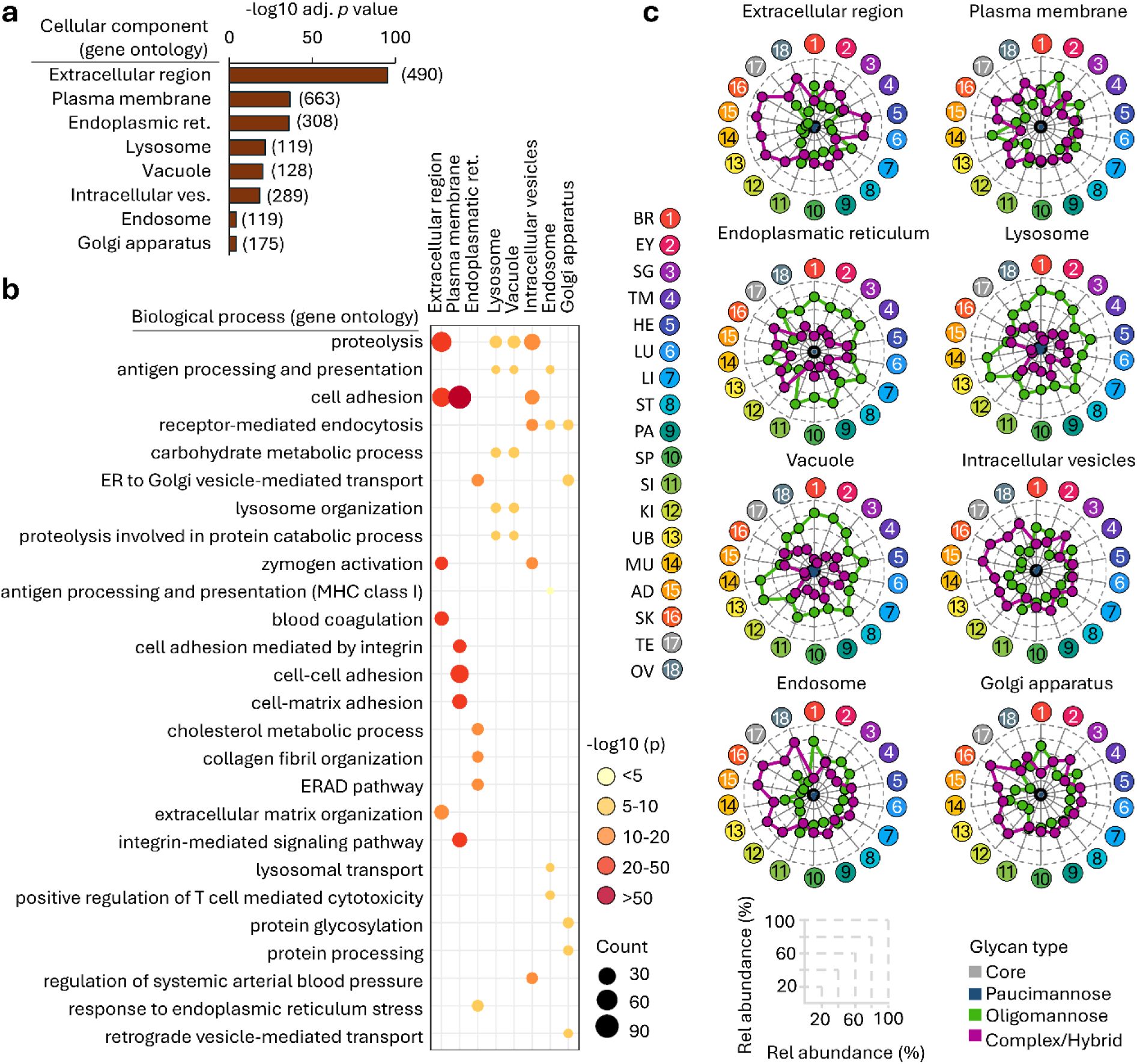
Distinct subcellular-specific *N*-glycosylation patterns and functional associations across tissues. **a**) Subcellular distribution of the global mouse *N*-glycoproteome (glycoproteins identified across all mouse tissues) as predicted by SubcellulaRVis online tool^60^. Only subcellular compartments that were enriched (adjusted *p* < 0.05) are shown with the number of enriched unique proteins indicated in brackets. **b**) Biological processes associated with the global mouse *N*-glycoproteome for glycoproteins mapped to distinct subcellular compartments. **c**) Glycoproteomics-based tissue-specific *N*-glycan type distribution by subcellular localization as annotated in panel a. See key and Supplementary Table S6 for details.

Discrete biological processes were associated with glycoproteins residing in different subcellular compartments. For example, proteolysis and cell adhesion were predominantly linked to proteins from the extracellular space and plasma membrane, respectively, supporting the known involvement of glycosylation in ECM remodeling and cellular interactions^25,26^ (**Figure 2b**). In contrast, antigen processing and presentation, as well as receptor-mediated endocytosis, were enriched among glycoproteins annotated to reside in intracellular compartments such as the lysosome, endosome, and the Golgi apparatus in agreement with current knowledge.

Consistent with previous reports^15,27,28^, glycoproteins mapped to the extracellular space, the plasma membrane and organelles carrying maturely processed glycoproteins (Golgi apparatus, endosomes, intracellular vesicles) were dominated by complex *N*-glycans (**Figure 2c**). However, even for this subset of glycoproteins, significant variations in glycan type distribution were observed across tissues, with oligomannose unexpectedly dominating in the brain, eye and within the small intestine (SI, #11). In contrast, and in line with literature^27^, glycoproteins annotated to reside in the ER, vacuoles and lysosomes displayed predominantly oligomannosidic-type glycans across most tissues with the exception of adipose tissue (AD, #15) and skin (SK, #16) exhibiting, for unknown reasons, relatively high levels of complex *N*-glycans across these intracellular compartments. Together, these findings document distinct compartment-specific glycosylation that vary dramatically across the mouse tissues and point to specialized tissue-specific functional roles of complex and oligomannosidic glycans that are tightly linked to the cellular localization and trafficking of their protein carriers.

### Variations in the glycosylation machinery drive glycoproteome diversity across mouse tissues

Next, we sought to explore mechanisms underpinning the tissue-specific glycoproteome diversity by correlating *N*-glycosylation features with the mRNA expression of a wide range of glyco-enzymes across key steps in the pathway for *N*-glycoprotein biosynthesis focusing on oligomannose processing and antennary branching, sialylation and fucosylation (**Figure 3a**). Glycoproteomics data revealed pronounced tissue-specific variation in oligomannose processing, with thymus exhibiting substantial trimming (36.9% M5 relative to precursor oligomannose species M6–M10), whereas minimal processing was observed in other tissues such as heart (HE, #5; 3.8%) and liver (LI, #7; 4.1%). (**Figure 3b**). This suggests that tissue-specific glycosylation arises, in part, from variations in the early ER-based maturation of nascent *N*-glycoproteins trafficking the secretory pathway. Supporting that late-stage glycan processing also contributes to tissue glycoproteome diversity, prominent inter-tissue differences were also observed for sialylation and fucosylation whereas comparable less variation were detected for antennary branching across the mouse tissues.

**Figure 3.**
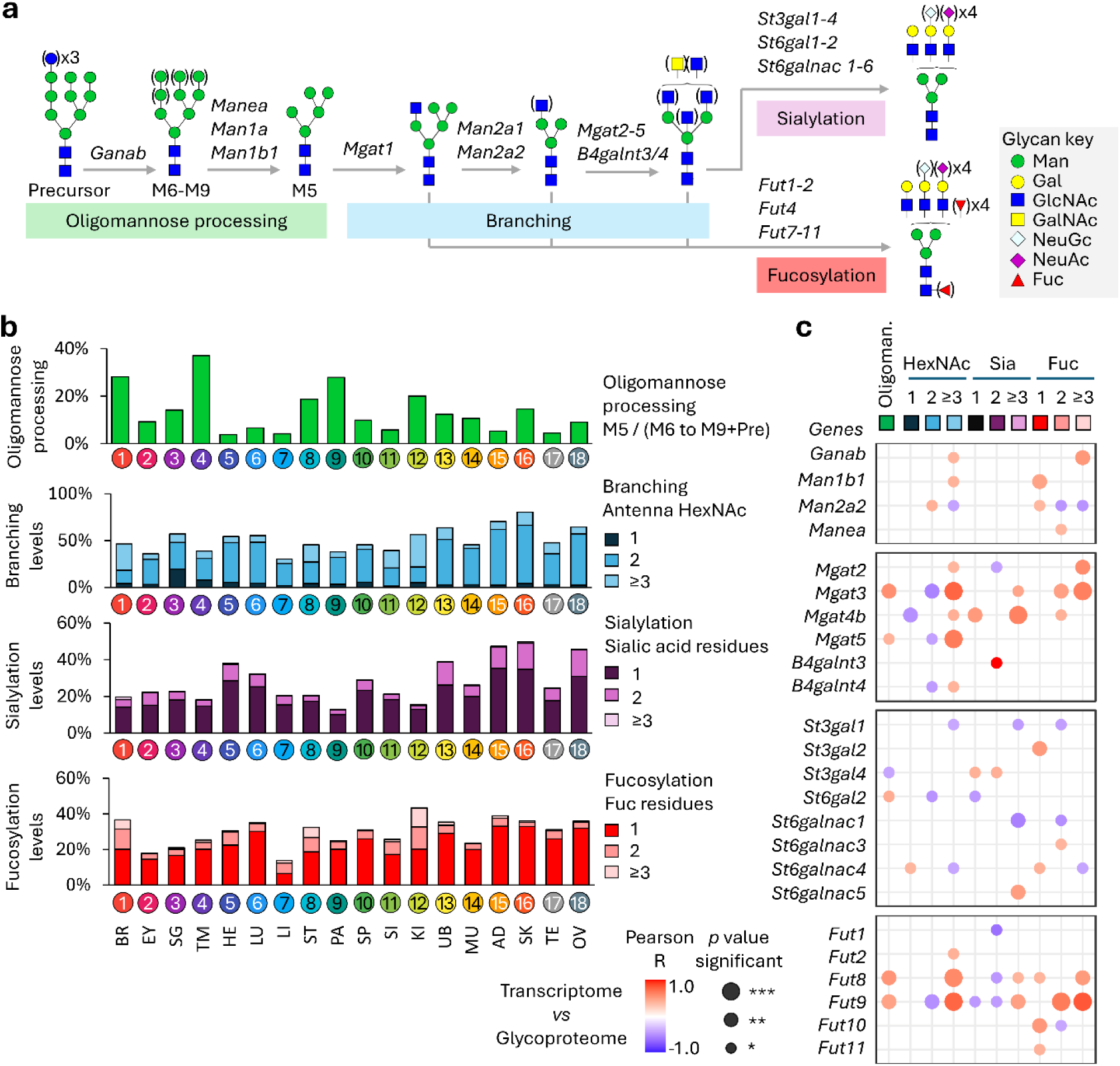
Tissue-specific *N*-glycosylation features and associations with glyco-enzyme expression patterns. **a)** Major steps and key glyco-enzymes responsible for the biosynthetic processing of *N*-glycoproteins. Some glycan-processing enzymes are left out for simplicity. **b)** Tissue-resolved *N*-glycan features quantified for the early ER maturation (oligomannose processing) and the downstream Golgi-based elongation stages involving antennary branching, sialylation and fucosylation as measured by LC-MS/MS glycoproteomics. **c)** Correlation between the global (combined across tissues) expression of glyco-enzymes (transcriptomics) and *N*-glycoproteome features (glycoproteomics). Only significant correlations are shown (Pearson R; \**p* < 0.05, \*\**p* < 0.01, \*\*\**p* < 0.001). See key for details.

Within the multi-omics framework, LMA analysis provided insight into a complementary glycome layer that enabled linkage- and epitope-sensitive interpretation of the glycoproteomics data^29^. Concordance was observed between the global oligomannose levels (combined across tissues) and the reactivity of mannose-reactive lectins (ConA, GNA, UDA, NPA) in the LMA data which was further supported by strong negative associations between these lectins and the level of complex-type *N*-glycans determined by glycoproteomics (**Supplementary Figure S2**). Fucosylation levels were also positively correlated with fucose-reactive lectins, including PSA and LCA, which recognize core fucosylation (α1,6-fucose), as well as AOL and AAL, which recognize diverse fucosyl linkages (α1,2-, α1,3-, α1,4-, α1,6-). Notably, sialylation was correlated with SNA, SSA, and TJA-I, which recognize α2,6-sialyl linkages, but not with MAL-I recognizing α2,3-sialylation.

Together, these correlations indicate that LMA profiling provides structurally informative glycome insights and support the view that the *N*-glycoproteome reflects the broader glycome architecture across mouse tissues, including other glycoconjugate classes such as *O*-glycans, glycosaminoglycans, and glycosphingolipids.

To explore possible associations to the glycosylation machinery, the global *N*-glycoproteome traits were assessed against the mRNA expression levels of relevant glyco-enzymes (**Figure 3c**). Strong positive associations were observed between highly branched *N*-glycans (≥3 antennae GlcNAc) and transcript levels of the Mgat family members, including *Mgat2*, *Mgat3*, *Mgat4b*, and *Mgat5*, as well as between antenna fucosylation (≥2 fucose residues) and *Fut8* and *Fut9*. In contrast, sialylation showed comparatively weaker associations to sialyltransferase expression patterns, with only *St3gal4* and *St6galnac5* correlating with mono- and di-sialylated glycopeptides. No associations were observed between oligomannose processing and the mannose-trimming enzymes (e.g. *Man1b1, Man2a2*). However, a considerable number of unexpected positive and negative correlations were identified between the glyco-enzymes and seemingly unrelated glycosylation features. For example, *St3gal1* and *St6galnac1* expression showed negative correlations with multi-sialylated glycans (≥3 sialic acid residues), while *Fut8* and *Fut9* were negatively and positively associated with di- and multi-sialylated glycans, respectively. These unexpected relationships suggest the presence of additional regulatory layers within the glycosylation machinery that remain to be elucidated in mouse tissues. Our data therefore provides an important resource to guide future efforts in systems glycobiology aiming to determine key glyco-enzymes and rate limiting steps in the glycosylation machinery that ultimately shapes the glycophenotypes exhibited by individual tissues.

### Diversity of sialic acid species contributes to tissue-specific sialylation

The mouse tissue glycome contains distinct sialic acid species, primarily *N*-acetylneuraminic acid (Neu5Ac) and *N*-glycolylneuraminic acid (Neu5Gc), which differ by a single hydroxyl group at the C5 position (**Figure 4a**). Using quantitative (HPLC-based) sialic acid profiling, we found that Neu5Ac and Neu5Gc are expressed globally in mice, but in levels that vary dramatically between tissues (**Figure 4b** and **Supplementary Table S5**). Brain and salivary gland (SG, #3) were highly enriched in Neu5Ac (both >90% of all sialic acid species), whereas liver and skin showed higher levels of Neu5Gc (all >75% of all sialic acid species). The modified sialic acids, such as *O*-acetylated derivatives, were generally low abundant or negligible in the mouse tissues with modest levels detected in select tissues, including brain, eyes, salivary gland, adipose tissue, and skin. The less common 2-keto-3-deoxy-D-glycero-D-galacto-nononic acid (Kdn) was detected only in the salivary gland. Levels of Neu5Gc and Neu5Ac on *N*-glycoproteins measured by glycoproteomics recapitulated the HPLC-based measurements of sialylation carried by all glycoconjugates (**Figure 4c**), supporting that the *N*-glycoproteome contributes significantly to the mouse tissue sialome. Sialic acid biosynthesis proceeds through a series of enzymatic steps converting UDP-GlcNAc to CMP-Neu5Ac, a nucleotide-sugar donor required for Neu5Ac to be transferred to glycoproteins by various sialyltransferases^30^ (**Figure 4d**). In mice, CMP-Neu5Ac can be further hydroxylated to CMP-Neu5Gc by Cmah, introducing an additional layer of complexity to the tissue glycome. Sialic acids can also be removed from glycoconjugates by sialidases (neuraminidases), creating a regulatory enzymatic network that contributes to the dynamic tissue-specific sialome^31^.

**Figure 4.**
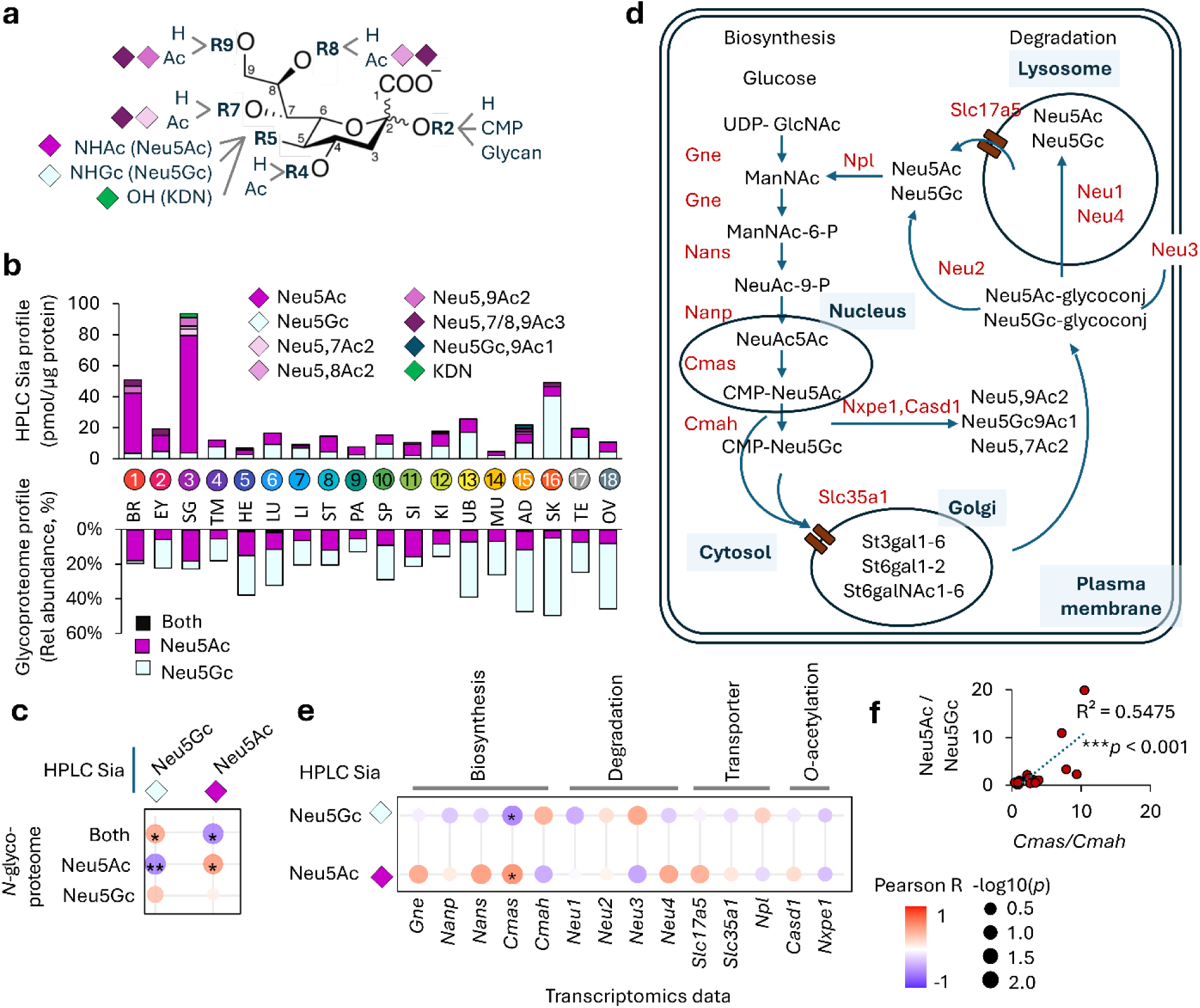
The mouse tissue sialome. **a)** Sialic acid species and modifications detected in mouse tissues. **b)** HPLC-based quantitation of the global tissue sialome (all sialic acid species across all tissues combined) (bars). Sialylation was also determined for the global *N*-glycoproteome as measured by LC-MS/MS glycoproteomics (mirror bars). **c)** Correlation between global sialic acid levels and sialylation of the tissue *N*-glycoproteome. **d**) Pathway overview of key substrates and enzymes involved in sialic acid biosynthesis and degradation. **e)** Correlation between tissue expression of enzymes mediating sialic acid anabolism and catabolism (highlighted in red in panel d) and sialylation levels as measured by HPLC profiling. For c and e, correlations were determined (Pearson R; \**p* < 0.05, ** *p* < 0.01). **f)** Correlation between global tissue levels of Neu5Ac/Neu5Gc as measured by HPLC and *Cmas*/*Cmah* gene expression as measured by transcriptomics.

Amongst several other less intuitive relationships, interrogation of the sialic acid profiles against the transcriptomics data revealed that the global levels of Neu5Ac and Neu5Gc (combined across tissues) correlate strongly with *Cmas* and *Cmah* expression (*p* < 0.001) (**Figure 4e-f**). While the many associations observed across these complex datasets highlight that much is yet to be understood in terms of enzymes that are directly and indirectly responsible for regulating the tissue sialome, the analyses point to two sialic acid precursor synthesis glyco-enzymes, Cmas and Cmah, being central in defining tissue-specific sialylation on *N*-glycoproteins^32,33^.

### Sex-specific N-glycoproteome traits across tissues

Sex differences in the *N*-glycoproteome across mouse tissues remain inadequately characterized generating an important knowledge gap that hampers both basic and applied research efforts in sex-biased glycobiology. Principal component analysis (PCA) using the full glycoproteomics datasets condensed to glycan compositional information illustrated distinct separation of males and females in a subset of tissues (**Figure 5**). Notably, the salivary gland, skin, kidney (KI, #12), heart, liver, and muscle were found to be sex-stratified tissues showing distinct separation between males and females. In contrast, the brain, adipose tissue, urinary bladder (UB, #13), lung (LU, #6), small intestine, and pancreas (PA, #9) showed no or negligible sex separation, while the thymus, spleen (SP, #10), stomach (ST, #8), and eye displayed only modest sex-specific differences when measured at the global glycan compositional level.

**Figure 5.**
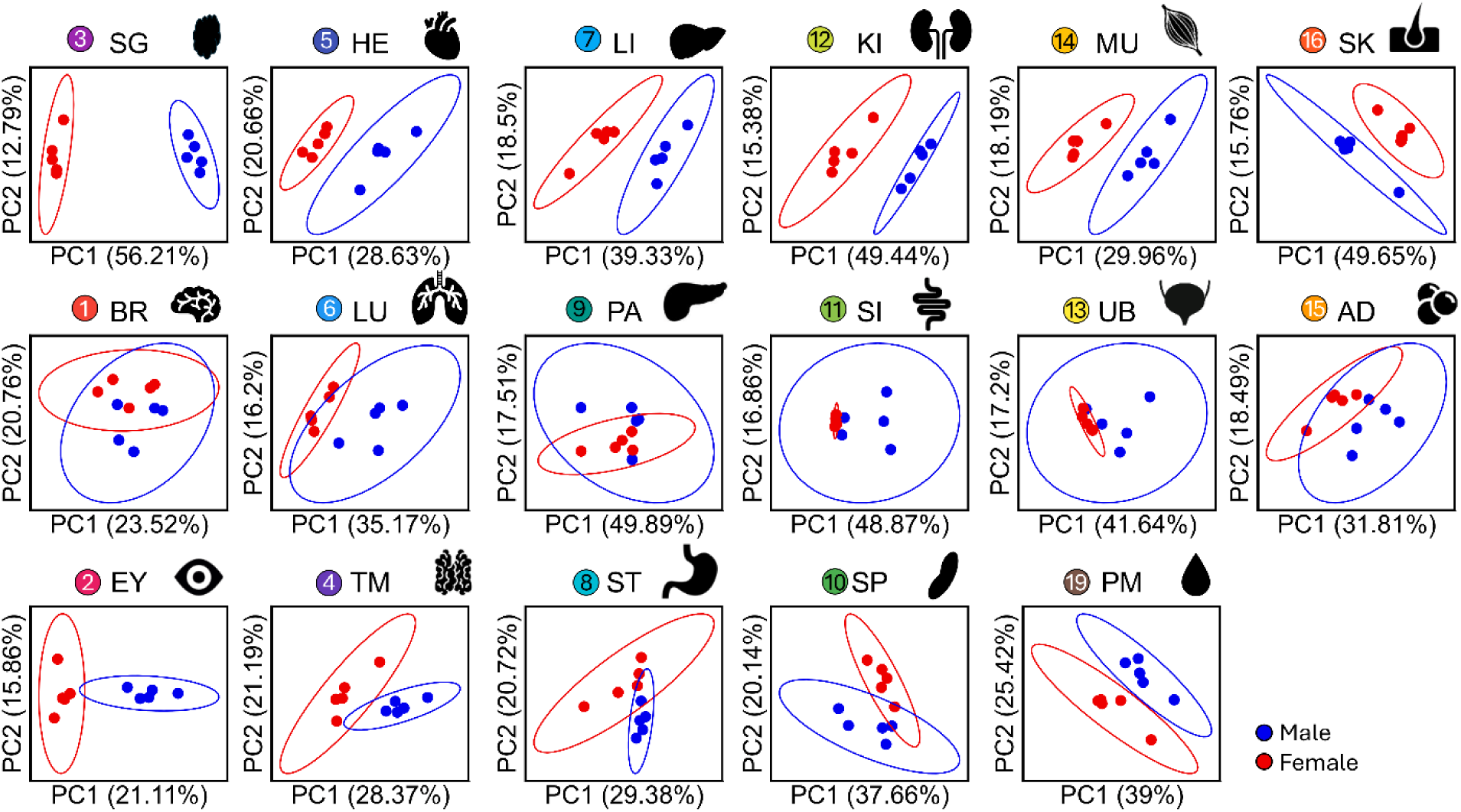
The sex-linked glycoproteome stratifies a subset of male and female mouse tissues. PCA plots were performed for each tissue using condensed glycan compositions extracted from the entire *N*-glycoproteomics datasets with males (blue) and females (red) separating in a subset of the investigated murine tissues. Sex-specific tissues (OV and TE) were left out of this analysis.

Prompted by these exciting findings, deeper mining of the glycoproteome data revealed profound sex-specific differences in oligomannose processing and antennary branching, sialylation and fucosylation across the mouse tissues (**Figure 6a**, see **Supplementary Table S6** for data). The salivary gland and liver, in particular, exhibited strong sex differences across all assessed glycosylation features, whereas the sex differences for most other tissues were isolated to one or two glycosylation traits. For example, muscle exhibited a higher level of antennary branching in males, and hyper-sialylation was a feature of the small intestines for females whereas the urinary bladder (UB, #13) displayed a strong bias for antennary fucosylation in male mice. Furthermore, Neu5Ac-type sialylation on *N*-glycoproteins was higher across a range of female tissues (SG, KI, SK, LU, MU, UB, EY, SP) whereas Neu5Gc levels were lower in the female brain, lung, thymus and stomach tissues (**Supplementary Figure S3a**). These findings were corroborated by HPLC sialic acid profiling, which confirmed elevated Neu5Ac in the female salivary gland and reduced Neu5Gc in female thymus and stomach tissues, highlighting sex-specific shifts in the sialome in select tissues (**Supplementary Figure S3b**).

**Figure 6.**
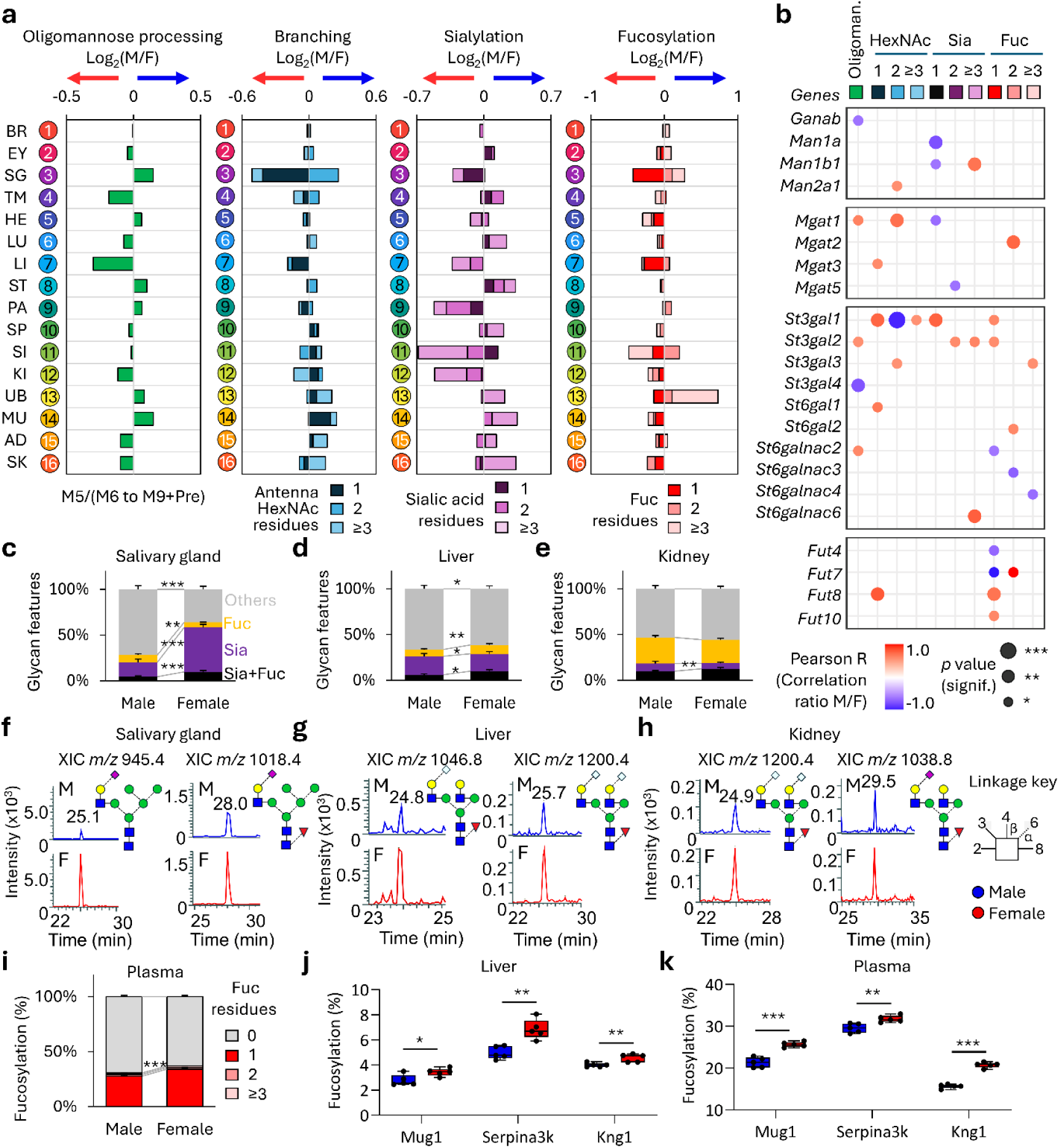
Sex-specific *N*-glycoproteome traits and glyco-enzyme expression patterns across tissues. **a**) Tissue-resolved profiling of glycan features in male and female mice as measured by LC-MS/MS-based glycoproteomics. Sex differences are expressed as log₂ fold change (male vs female). **b**) Correlation between sex differences in the global *N*-glycoproteome and the expression patterns of relevant glyco-enzymes (transcriptomics). Only significant correlations are shown, see key for details (R; \**p* < 0.05, \*\**p* < 0.01, \*\*\**p* < 0.001). Sex-specific differences in fucosylation and sialylation as determined by PGC-LC–MS/MS *N*-glycomics for the salivary gland, liver, and kidney shown as the proportion of the tissue *N*-glycome (**c-e**) and plotted for representative core fucosylated and α2,6-sialylated *N*-glycan structures for males (top) and females (bottom) (**f-h**). **i**) Plasma fucosylation profile in male and female mice measured by glycoproteomics. Fucosylation levels of Mug1, Serpina3k, and Kng1 in **j**) liver and **k**) plasma. For all sex comparisons, unpaired two-tailed t-test; \**p* < 0.05, \*\**p* < 0.01, \*\*\**p* < 0.001. See Figure 3 for glycan key.

Integration of the sex-stratified global glycoproteome features (combined across tissues) with the glyco-enzyme expression patterns across male and female mice yielded clues to the molecular mechanisms that may underlie the sex-linked glycophenotypes. Sex differences in antennary branching, sialylation, and fucosylation correlated positively with altered mRNA expression of *Mgat1*, *St3gal1*, *St3gal2*, *Fut7*, *Fut8*, and *Fut10* in male vs female mouse tissues pointing to glyco-enzymes that may contribute to the sex-specific glycosylation observed in some tissues (**Figure 6b**).

Quantitative proteomics showed that the proteome remained largely stable between male and female tissues (**Supplementary Figure S4a-b** and **Supplementary Table S7** for data), suggesting that other molecular traits are the main contributors to sex differences in most tissues. However, pronounced sex-dependent proteome differences were observed in the salivary gland, liver, and kidney, which were also accompanied by strong differences in the glycoproteome (**Supplementary Table S8**). Notably, the salivary gland exhibited the most pronounced sex-specific variation (over 50%) in both the glycoproteome and proteome, consistent with the distinct protein band patterns observed between male and female salivary gland tissues (**Supplementary Figure S4b**). In contrast, skin and plasma (PM, #19) showed prominent sex-dependent differences at the glycoproteome level despite no detectable differences in the overall proteome between males and females.

To further explore the profound sex-specific glycophenotypes of the salivary gland, liver, and kidney at the detailed glycome level, *N*-glycans were released from these tissues and profiled by PGC-LC-MS/MS with a focus on sialylation and fucosylation (**Supplementary Table S9**). Consistent with the glycoproteomics data, glycans carrying sialic acid and fucose were elevated in the *N*-glycome of the female salivary gland, liver, and kidney (**Figure 6c-e**) as also documented for individual sialylated and fucosylated *N*-glycans across those three tissues (**Figure 6f-h**). Deep structural analysis of *N*-glycans from the salivary gland, liver, and kidney by PGC-LC-MS/MS, together with correlations to fucose-reactive PSA and LCA lectins, confirmed that the increased fucosylation in female tissues was driven by elevated α1,6-fucosylation (core fucose) while α2,6-sialylation was found to dominate these three sex-specific tissues (**Supplementary Figure S5**). These data indicate that lectin reactivity patterns observed in the LMA analysis were consistent with the linkage-specific glycan structures identified by PGC-LC–MS/MS, particularly the enrichment of α1,6-fucosylation and α2,6-sialylation in sex-biased tissues.

Importantly, the increased fucosylation observed in female liver was also mirrored in female plasma (**Figure 6i**). This observation is consistent with the liver being the primary source of circulating plasma glycoproteins^34^ and suggests that sex-biases in hepatic glycosylation is reflected systemically. In support, increased fucosylation of female liver and plasma were observed for multiple liver-derived plasma proteins including murinoglobulin (Mug1), serine protease inhibitor A3K (Serpina3k), and kininogen-1 (Kng1) (**Figure 6j-k**).

### Glyco-enzymes driving sex-specific glycosylation in mouse tissues

To further explore mechanisms driving sex-linked glycoproteome variations in mouse tissues, we integrated glycoproteomics, proteomics, and transcriptomics to determine if the observed sex differences were driven primarily by protein-specific glycoform alterations, by changes in overall protein abundance, or by variations in glyco-enzyme expression across male and female tissues. In the salivary gland, the many glycoforms that differ between males and females (x-axis) were accompanied by sex-based alterations in carrier protein abundance (y-axis) (**Figure 7a**). Sex-specific sialylation and fucosylation patterns were observed across multiple proteins some of which exhibited strong protein level alterations and others that appeared altered independently of any protein-level changes, as exemplified by Muc19 and Tmed9, respectively. The many sex-linked glycoproteome differences in the salivary gland were underpinned by strong sex-specific expression patterns of glyco-enzymes including several glycosyltransferases (*Mgat1, St3gal1, St3gal4, St6galnac2, Fut2, Fut8*) and genes related to sialic acid and fucose biosynthesis (*Cmah, Tsta3, Slc17a5, Slc35a1*) that may directly or indirectly influence the discrete sex phenotypes of the male and female salivary gland glycoproteome.

**Figure 7.**
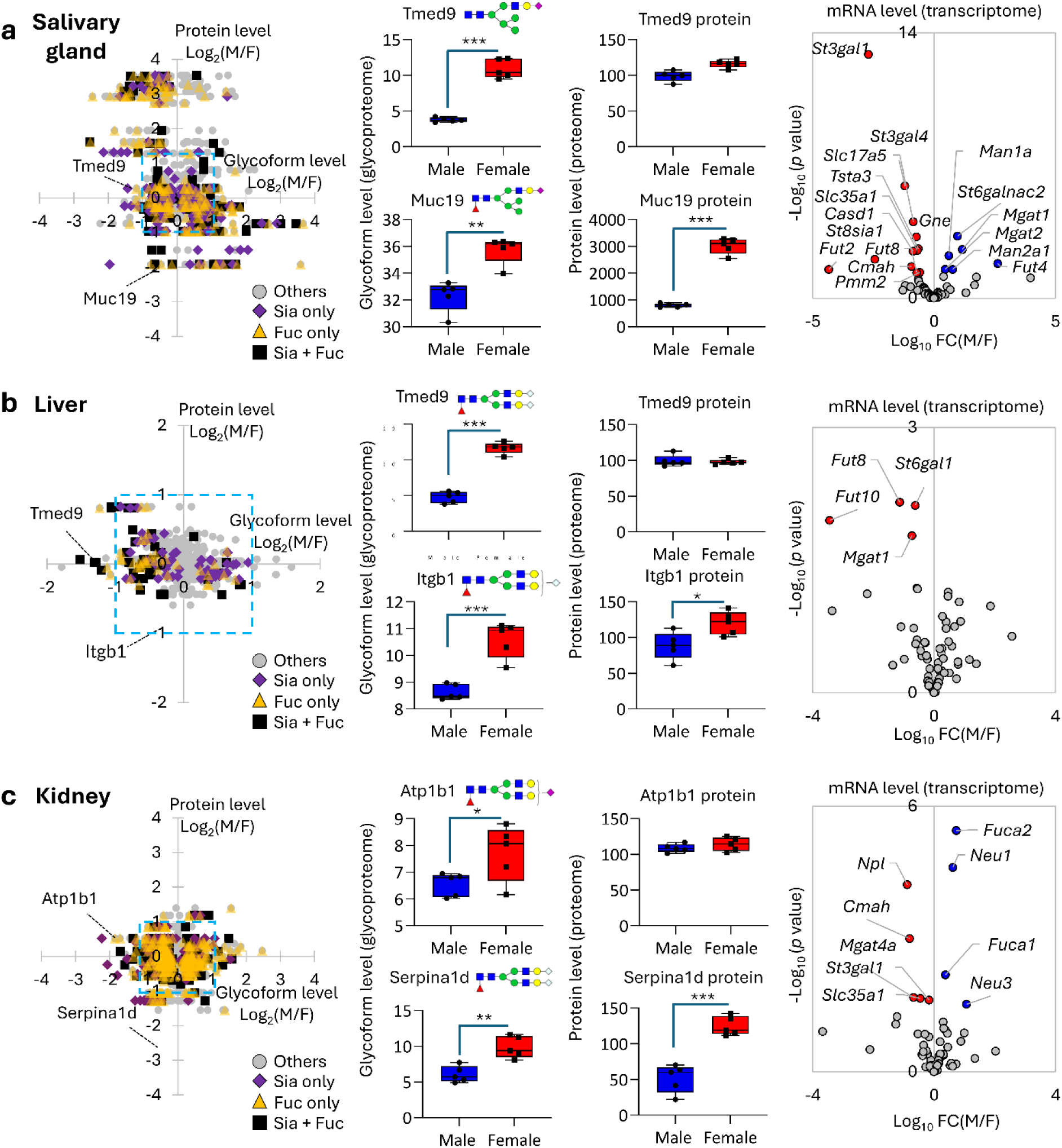
Mechanistic clues to the sex-specific glycophenotypes of mouse salivary gland, liver and kidney. Sex differences in the *N*-glycoproteome (glycoproteomics, x-axis) plotted against sex differences in the proteome (proteomics) in the **a)** salivary gland, **b)** liver, and **c)** kidney (left). Sex-specific differences in sialylation (purple diamonds), fucosylation (yellow triangles), and sialo-fucosylation (black squares) are highlighted (t-test, FDR < 0.05). Data points outside the blue confidence box represent proteins with an absolute fold change > 2 between males and females. Relative levels of select glycoforms and proteins that show prominent sex differences are plotted (middle). Volcano plots comparing the glyco-enzyme expression profiles in males and females (right). Transcripts elevated in females are in red whereas transcript elevated in males are in blue (glmQLFTest, *p* < 0.05). See Figure 3 for glycan key.

The liver displayed comparatively fewer sex differences at the individual glycoprotein level (**Figure 7b**). Of the altered glycoproteins, females exhibited broadly increased sialylation and fucosylation (x-axis), largely independent of changes in overall protein abundance (y-axis). Similar to the salivary gland, sialofucosylated Tmed9 glycoforms, but not Tmed9 protein levels, were higher in female livers whereas the elevation of sialofucosylated Itgb1 glycoforms in female livers, similar to salivary gland Muc19 was accompanied by protein level alterations. Notably, a significant increase in the transcript levels of sialyltransferase *St6gal1* and the fucosyltransferases *Fut8* and *Fut10* was observed in females, suggesting that relatively restricted male and female differences in the glycosylation machinery contribute to the sex-specific glycoproteome in the liver.

In kidney, extensive variations in sialylated and fucosylated glycoforms were observed across males and females, but without strong accompanying sex-linked differences in protein abundance (**Figure 7c**). The transcript levels of *St3gal1* and enzymes related to sialic acid synthesis (*Slc35a1, Cmah*) were significantly elevated in females, while fucosidases (*Fuca1, Fuca2*) and neuraminidases (*Neu1, Neu3*) were higher in males suggesting a complex interplay of competing glycan synthesis and degradation pathways underpinning the sex-specific glycoproteome of the kidney.

To provide a comprehensive resource for exploring sex-linked glycosylation, we developed an interactive online database that enables interrogation of the *N*-glycoproteome at the individual protein and site level across male and female mouse tissues (https://igcore.cloud/mta/atlas-viewer/). The platform enables users to compare glycosylation at multiple levels, from broad glycan classes, including antennary branching, sialylation, and fucosylation, to specific glycoforms, alongside corresponding protein abundance information. The platform allows sex comparisons across all tissues or selected tissue subsets, offering a detailed view of sex variations in the mouse tissue glycoproteome. This online resource serves as a tool for researchers to dissect the sex-specific glycoproteome differences, obtain clues to their molecular mechanisms and discover links to sex-biased tissue functions and diseases.

A summary visualizing the multiple molecular layers of the sex-specific glycophenotypes observed across the mouse tissues is presented in **Figure 8**, highlighting salivary gland, liver, and kidney as the three strongest sex-linked tissues in terms of *N*-glycosylation, whereas eye, brain, and spleen exhibited the lowest degree of variation across biological sex.

**Figure 8.**
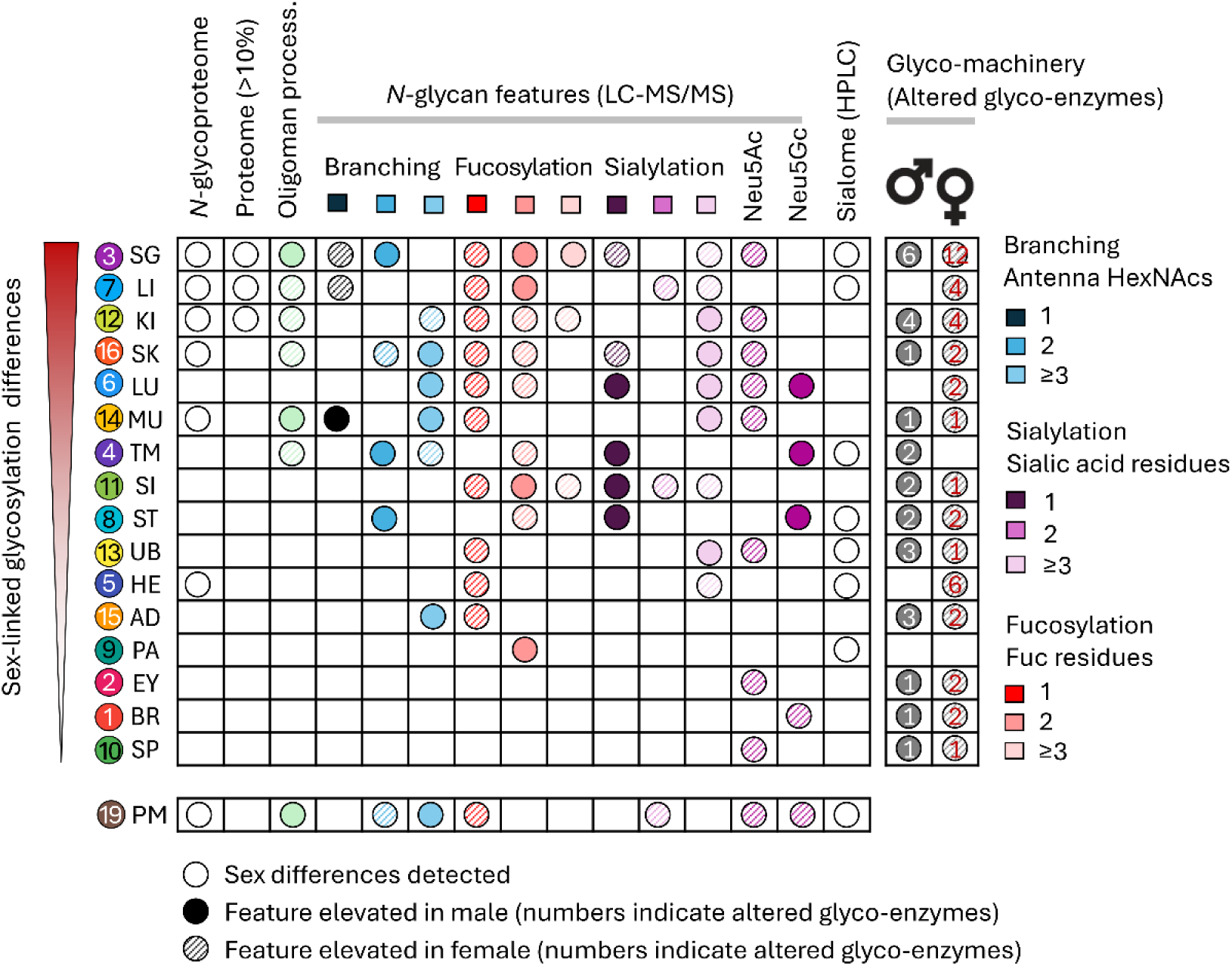
The sex-specific glycophenotypes of murine tissues. Schematic overview integrating proteomics, glycoproteomics, glycomics, and transcriptomics findings to illustrate the molecular basis of sex-linked glycosylation differences across mouse tissues. See key for details.

## Discussion

Our multi-omics study reveals that biological sex exerts a profound influence on tissue glycosylation, with the salivary gland, liver, and kidney showing pronounced sex-linked glycoproteome variations, whereas the brain maintains extensive glycome diversity that is largely conserved between male and female mice. These findings highlight that sex influences glycosylation in a tissue-dependent manner, reflecting coordinated regulation of the glycosylation machinery and protein expression according to organ-specific physiological requirements.

Increased core fucosylation in females was consistently observed across multiple tissues, including the salivary gland, liver, kidney, skin, lung, muscle, small intestine, urinary bladder, heart, and adipose tissue, observations that unsurprisingly correlated with transcript levels of *Fut8* in those tissues. The liver, as the primary source of plasma proteins, displayed pronounced sex-specific fucosylation patterns, which was reflected in several liver-derived plasma glycoproteins (e.g. Mug1, Serpina3k, Kng1). These observations are consistent with previous reports showing higher core fucosylation in plasma from female mice^21,35^ and, importantly, also in human plasma^19,20,22^, indicating that glycophenotypic observations made in this mouse model may map to other mammals and guide functional and translational efforts aiming to understand human sex-biased tissue biology. Importantly, these findings also highlight the relevance of sex as a critical variable in glycan biomarker discovery. As many clinically investigated glycobiomarker candiates are liver-derived plasma glycoproteins, inherent sex-dependent differences in glycosylation, such as the elevated core fucosylation observed in females, may influence baseline biomarker levels and their interpretation. Consequently, incorporating sex-stratified analyses in glycobiomarker research will be essential to improve biomarker specificity, sensitivity, and translational applicability across populations.

The liver is highly responsive to sex hormone regulation through estrogen and androgen receptors, which modulate gene expression and enzymatic activity^36^. While our knowledge of how sex hormones impact hepatic glyco-enzyme expression is only just emerging^37^, it has been established that core fucosylation plays critical roles in liver physiology by influencing protein folding, stability, and secretion of key plasma glycoproteins, including coagulation factors, transport proteins, and acute-phase reactants^38^. Altered core fucosylation, as observed across male and female livers, can therefore affect hepatic function, impacting processes such as protein clearance, receptor-mediated signaling, and intercellular communication. Notably, changes in α1,6-fucosylation have also been linked to liver repair and regeneration^39^, as well as susceptibility to liver diseases, including fibrosis^40^, steatosis^41^, and hepatocellular carcinoma^42^. These findings suggest that sex-linked differences in Fut8-mediated core fucosylation may play roles in the physiological differences between male and female livers thereby potentially contributing to generating sex-biased liver disease risk as for example observed in patients with liver cirrhosis^43^. Sex-specific differences in sialylation were most pronounced in the salivary gland, where increased sialylation, predominantly Neu5Ac species, was consistently detected in female tissues by both glycoproteomics and HPLC-based sialic acid profiling. In female salivary glands, protein-and site-specific elevation of sialylated glycoforms, predominantly α2,6-sialylation, were observed across a broad range of glycoproteins, including secreted components such as mucin-19. Sialic acids play critical roles for the function of salivary glycoproteins by mediating mucosal lubrication, regulating microbial adhesion, and providing protection against viral and bacterial pathogens^44–46^. Therefore, sex differences in sialylation within the salivary gland are likely to influence oral immune defense, microbiome composition, and susceptibility to infection as well as impacting the physicochemical properties of saliva that support tissue homeostasis and barrier function. However, future studies are required to determine if sialylation or any other glycosylation features are directly or indirectly related to sex-biased disorders of the salivary gland such Sjögren’s (dry mouth) syndrome that overwhelmingly impacts women^47^.

In contrast to peripheral tissues, the brain displayed remarkably conserved glycosylation between males and females, with no major sex differences detected within the *N*-glycoproteome. This molecular stability was paralleled by the absence of major sex-linked alterations in both the proteome and transcriptome, indicating that the biosynthetic machinery governing neural glycosylation is tightly regulated and largely sex-independent under physiological conditions. In line with recent literature^35,48–50^, the brain *N*-glycome instead exhibited extensive molecular diversity through well documented structural features, including high glycan complexity and extensive terminal modifications, consistent with the specialized requirements of neural tissue for synaptic function, cell–cell communication, and receptor signaling. The concordance across the glycoproteome, proteome, and transcriptome layers suggests that, unlike metabolically and hormonally responsive organs, the brain maintains a buffered glycosylation program that is resilient to systemic sex hormone fluctuations. Functionally, this conservation likely reflects evolutionary pressure to preserve neural circuitry and cognitive processes, where even subtle molecular perturbations could have profound physiological consequences. Collectively, these findings position brain glycosylation as a stable molecular framework, highlighting tissue-specific vulnerability or resistance to sex-dependent regulation across the organism.

This study has several limitations that should be considered when interpreting the findings. First, analyses were conducted at the bulk tissue level, which masks cellular heterogeneity and precludes resolution of sex-specific glycosylation differences within specific cell types or micro-environmental niches. Second, although translational parallels were observed, inherent species barriers exist between mice and humans, including variations in sialic acid composition most notably the absence of human Neu5Gc and differences in glycosyltransferase repertoires, requiring cautious extrapolation of these results to human biology. Finally, the study captures glycosylation landscapes at a single age time point and for a single mouse strain (C57BL/6J) only investigated under basal physiological laboratory conditions, thereby not accounting for dynamic regulation across development, aging, hormonal transitions, genotypic diversity or pathophysiological conditions. Collectively, these limitations highlight the need for future investigations incorporating single-cell resolution, cross-species validation, and longitudinal designs in relevant disease models to further refine our understanding of sex-linked glycophenotypes. Additionally, we acknowledge that this the *N*-glycoproteome-focused study deliberately overlooks the many other glycoconjugates that also exist within the vast tissue glycome and that similarly may contribute to sex-biased tissue functions.

Nevertheless, this study moves beyond descriptive glycome mapping to establish a mechanistic framework linking glycan structures to their regulatory networks across tissues, revealing how biological sex shapes glycosylation through coordinated control of enzymatic pathways and substrate protein and nucleotide-sugar availability. Rather than capturing global glycome shifts alone, we integrate comparative glycoproteomics, proteomics, and transcriptomics with detailed structure-focused glycomics to resolve sex-dependent regulation across three interconnected layers: site-specific glycan distribution on defined protein carriers, the abundance of those carriers, and the activity of the glycosylation biosynthetic machinery. Such system-level resolution has not been achieved in prior body-wide tissue studies, which have largely focused on glycosylation features without resolving protein-site specificity, protein level regulation and glycosylation machinery status.

To maximize accessibility and translational utility of our findings, we provide an interactive online viewer (https://igcore.cloud/mta/atlas-viewer/) that enables exploration of glycosylation in male and female tissues at the protein- and site-specific level. This resource constitutes a foundational platform for hypothesis generation and investigation of sex-biased disease mechanisms, collectively advancing our knowledge of the molecular features and functions of the sex-linked glycoproteome.

## Methods

### Animals care and ethics statement

C57BL/6J mice (five males, five females, all 8 weeks of age) were obtained from Japan SLC (Shizuoka, Japan) and maintained in a controlled environment (23 ± 2°C, 50 ± 10% relative humidity, 12:12 light/dark cycle) for 2 weeks with food and water available *ad libitum*. At 10 weeks, the mice were euthanized under deep isoflurane anesthesia and individual tissues were collected. All procedures were approved by the Animal Care and Use Committee of Nagoya University (permit: A240095-003) and carried out in accordance with both the institutional and ARRIVE guidelines (https://arriveguidelines.org). Every effort was made to limit the number of animals used and minimize their suffering.

### Tissue and blood collection

Blood was collected from the right atrium immediately after sacrifice in EDTA tubes. Plasma (PM, #19) was separated by centrifugation at 1,000 x *g* for 10 min at 4°C and kept at -80°C until further handling. Subsequently, PBS perfusion was performed to remove residual blood. Brain (whole, BR, tissue #1), eyes (EY, #2), salivary gland (SG, #3), thymus (TM, #4), heart (HE, #5), lung (LU, #6), liver (LI, #7), stomach (ST, #8), pancreas (PA, #9), spleen (SP, #10), small intestine (SI, #11), kidney (KI, #12), urinary bladder (UB, #13), muscle (MU, #14), adipose tissue (AD, #15) and skin (SK, #16) were surgically removed by a trained animal technician. For the male mice, the testes (TE, #17) were also surgically removed as were the ovaries (OV, #18) from the female mice. Tissues were kept at -80°C until further handling.

### Protein preparation for glycoproteomics and proteomics

Frozen tissues were lysed and homogenized in 500 µl lysis buffer containing PBS, 1% (v/v) Triton, 1 mM EDTA, 1 mM PMSF, 10 mM sodium fluoride, 1 mM sodium orthovanadate and a protease inhibitor cocktail (Roche) using zirconium beads (3 mm diameter, Sigma) in a TissueLyser (30 Hz, 2 min, Qiagen). Protein concentrations were determined by BCA (Thermo) ahead of protein precipitation using four volumes of acetone (16 h, -30°C). Proteins were pelleted (14,000 rpm, 10 min, 4°C) and resuspended in 8 M urea in 50 mM TEAB.

Protein extracts from tissues were reduced using 10 mM DTT (30 min, 30°C) and alkylated using 40 mM iodoacetamide (final concentrations, 30 min, in the dark, 20°C). The alkylation reactions were quenched using excess DTT. Samples were digested using sequencing grade porcine trypsin (1:50, w/w; 12 h, 37°C, Promega). Proteolysis was stopped by acidification using 1% (v/v) trifluoroacetic acid (TFA, final concentration). Peptides were desalted using primed Oligo R3 reversed phase solid phase extraction (SPE) micro-columns and dried^51^.

### TMT labelling of peptides

Peptides were labelled with tandem mass tags (TMT) according to the manufacturer’s protocol. Separate peptide reference pools were generated by pooling 2 µg of the peptide mixture from all tissue samples to enable quantitative comparisons across multiple TMT-10plex experiments. The 131C reporter ion channel was consistently used as the reference channel.

For each tissue, peptides derived from 20 µg protein extracts of five male and five female samples were dissolved in 50 µl of 100 mM TEAB and subsequently labelled with TMT11plex reagents (0.16 mg in 20 µl anhydrous acetonitrile, 1 h at 20°C; Thermo). Labelling reactions were quenched using 4 μl 5% (v/v) hydroxylamine (15 min, 20°C). Labelled peptides were mixed 1:1 (w/w) and desalted using hydrophilic–lipophilic balance SPE cartridges (Waters). A small aliquot containing 10 µg peptide mixture was set aside for direct LC-MS/MS analysis (unenriched fraction) and dried. The remaining samples were dried for glycopeptide enrichment.

### Glycopeptide enrichment

TMT-labelled peptide mixtures (∼220 μg combined) were reconstituted in 50 μl 80% ACN in 1% (both v/v) TFA and loaded onto primed custom-made hydrophilic interaction liquid chromatography (HILIC) SPE micro-columns packed with zwitterionic ZIC-HILIC resin (10 μm particle size, 200 Å pore size, kindly provided by Merck Millipore) onto supporting C8 disks (Empore) in p10 pipette tips^51,52^. Briefly, the flow-through and wash fractions containing non-glycosylated peptides were combined and kept for separate downstream analysis. The retained glycopeptides were eluted sequentially with 0.1% (v/v) TFA, 25 mM ammonium bicarbonate, and then 50% (v/v) ACN. Eluted fractions were pooled. The unenriched and HILIC flowthrough (peptides) and enriched (glycopeptides) fractions were separately dried, desalted on primed Oligo R3 reversed phase SPE micro-columns (as above), aliquoted, and dried.

### LC-MS/MS

Unenriched peptide and enriched glycopeptide samples were separately injected on a PepMap™ Neo Nano Trap Cartridge (5 mm × 300 μm inner diameter, Thermo) and separated on an analytical LC column (Aurora Ultimate; 25 cm × 75 μm, 1.7 μm ID, IonOpticks) at 300 nl/min delivered by a Vanquish Neo UHPLC System (Thermo). The mobile phases were 99.9% ACN in 0.1% (both v/v) formic acid (FA, solvent B) and 0.1% (v/v) FA (solvent A). The gradient was 3-35% B over 90 min, 35-50% B over 8 min, 50-90% B over 2 min, and 10 min at 95% B. The nanoLC was connected to an Orbitrap Exploris 240 mass spectrometer (Thermo) operating in positive ion polarity mode.

For enriched glycopeptide fractions, the Orbitrap acquired full MS1 scans (*m/z* 500-2,000, AGC: standard, 100 ms maximum accumulation, 120,000 FWHM resolution at *m/z* 200). Employing data-dependent acquisition with a fixed 3 s cycle time, abundant precursor ions from each MS full scan were ranked and sequentially isolated and fragmented utilizing stepped higher-energy collision-induced dissociation (HCD) at normalized collision energies (NCEs) of 20%, 30%, and 40%. Only multicharged precursors (Z ≥ 2) were selected for fragmentation. Fragment spectra were acquired in the Orbitrap with the following settings: 45,000 resolution, AGC: standard, 200 ms maximum accumulation time, fixed first mass set to *m/z* 110, *m/z* 1.6 precursor isolation window and 20 s dynamic exclusion after a single isolation and fragmentation of a given precursor. For unenriched peptide fractions, the Orbitrap acquired full MS1 scans (*m/z* 380-1,800, AGC: standard, 100 ms maximum accumulation, 120,000 FWHM resolution at *m/z* 200). Employing data-dependent acquisition with a fixed 3 s cycle time, abundant precursor ions from each MS full scan were ranked and sequentially isolated and fragmented with HCD-MS/MS utilizing an NCE of 30%. Only multicharged precursors (Z ≥ 2) were selected for fragmentation. Fragment spectra were acquired in the Orbitrap with the settings: 45,000 resolution, AGC: standard, maximum accumulation time set to “auto”, fixed first mass set to *m/z* 110, *m/z* 1.4 precursor isolation window and 20 s dynamic exclusion after a single isolation and fragmentation of a given precursor.

### Glycoproteomics and proteomics data processing and analysis

For glycoproteomics, HCD-MS/MS data were searched with Byonic (v2.6.46, Protein Metrics) using 10/20 ppm as the precursor/product ion mass tolerance, respectively. Cys carbamidomethylation (+57.021 Da) and N-term/Lys TMT (+229.163 Da) were considered fixed modifications. Fully tryptic peptides were searched with up to two missed cleavages per peptide. For the variable modifications, up to two common and one rare modification(s) were permitted per peptide. Methionine oxidation (+15.994 Da) was defined as a common modification, while *N*-glycosylation of sequon-localized asparagine residues was specified as a rare modification. The *N*-glycan search space comprised 309 predefined mammalian structures available in Byonic (excluding sodium adducts) manually supplemented with an additional 151 glycomics-informed entries from a recent high-quality *N*-glycome map of mouse tissues^14^ for a total glycan search space of 460 glycan compositions. The HCD-MS/MS data were searched against the UniProt Reference *Mus musculus* proteome (17,150 reviewed sequences, UniProtKB, released June 2023). All searches were filtered to <1% false discovery rate (FDR) at the glycoprotein and glycopeptide level using a protein decoy database. Only confident glycopeptide identifications (PEP 2D < 0.01) were considered. Glycopeptides were quantified using the ‘Report Ion Quantifier’ available as a node in Proteome Discoverer (v2.2, Thermo)^52^. Glycopeptides were manually grouped by summing the reporter ion intensities from the glycopeptide-to-spectrum matches (glycoPSMs) belonging to the same UniProtKB identifier, same glycosylation site within each protein, and same glycan composition. Within each sample, the total set of glycopeptide forms from each glycosylation site from each protein were normalized to 100%.

For proteomics, HCD-MS/MS data (from both unenriched and HILIC flow-through fractions) were processed using Proteome Discoverer v3.0 and searched against UniProt Reference *Mus musculus* proteome (17,150 reviewed sequences, UniProtKB, released June 2023) using the Sequest-HT search engine with a mass tolerance of 5 ppm for precursor ions and 10 ppm for product ions. The enzyme specificity was set to trypsin with a maximum of two missed cleavages permitted. Carbamidomethylation of Cys (+57.021 Da) and N-term/Lys TMT (+229.163 Da) were fixed modifications whereas Met oxidation (+15.994 Da) was a variable modification. Both protein and peptide level identifications were filtered to 1% FDR. The TMT-based protein abundances were normalized by ratio to the reference channel and total peptide as previously described^18^.

### Glycan preparation for glycomics

In preparation for glycomics, protein extracts (15 µg/sample) from kidney, liver and salivary gland pooled from five male and five female mice were immobilized on a primed 0.45 μm polyvinylidene fluoride membrane (Merck Millipore) and handled as described^53^. Briefly, *N*-glycans were released using 10 U recombinant *Elizabethkingia miricola* peptide:*N*-glycosidase F (10 U/μl, 16 h, 37°C, Promega). Detached glycans were reduced using 1 M sodium borohydride in 50 mM potassium hydroxide (3 h, 50°C). Reactions were stopped using glacial acetic acid, and glycans were desalted using strong cation exchange/C18 and porous graphitized carbon (PGC) SPE micro-columns^54^.

### PGC-LC/MS-MS glycomics

Released *N*-glycan mixtures were separated on an UltiMate 3000 HPLC system (Dionex) interfaced with a LTQ Velos Pro linear ion trap mass spectrometer (Thermo). Glycans were loaded on a PGC HPLC capillary column (Hypercarb KAPPA, 5 μm particle size, 200 Å pore size, 180 μm inner diameter × 100 mm length, Thermo) operated at 50°C with a constant flow rate of 20 μl/min. The mobile phases were 10 mM ammonium bicarbonate, pH 8.0 (solvent A) and 10 mM ammonium bicarbonate in 70% (v/v) ACN (solvent B) employing a 86 min gradient: 8 min at 2.6% B, 2.6-13.5% B over 2 min, 13.5-37.3% B over 55 min, 37-64% B over 10 min, 64-98% B over 1 min, 5 min at 98% B, 98-2.6% B over 1 min, and 4 min at 2.6% B. The electrospray ionization source was operated in negative ion polarity mode with a source potential of 3.6 kV. Full MS1 scans (*m/z* 570-2,000) were acquired using one micro-scan, *m/z* 0.25 FWHM resolution, 5 × 10^4^ AGC, and 50 ms maximum accumulation time. MS/MS data were acquired using *m/z* 0.35 FWHM resolution, 2 × 10^4^ AGC, 300 ms maximum accumulation time, and 2 *m/z* precursor ion isolation window. The five most abundant precursors from each MS1 full scan were selected for collision-induced dissociation–based MS/MS using a NCE of 33% with an activation Q of 0.250 and 10 ms activation time.

### Glycomics data analysis

Xcalibur (v2.2, Thermo) was used to inspect, browse and annotate the raw LC–MS/MS data. Glycans were manually identified and quantified as described^52,55,56^. Briefly, glycan precursor ions were extracted using RawMeat (v2.1, Vast Scientific), common contaminants and redundant precursors removed, and generic monosaccharide compositions considering only Hex, HexNAc, dHex, Neu5Ac, Neu5Gc were established using GlycoMod (http://www.expasy.ch/tools/glycomod). The glycan fine structures were manually elucidated using monoisotopic precursor mass, PGC-LC elution time and MS/MS fragmentation pattern (using Glycoworkbench v2.1). Glycans were quantified using extracted ion chromatograms in Skyline (v9.1)^56^.

### Sialic acid speciation and profiling

Mouse tissue lysates were analyzed in technical triplicates. Lysates were hydrolyzed in 2 M propionic acid (Wako, Japan) at 80°C for 2 h, followed by drying. Dried samples were resuspended in 20 µl 0.01 M TFA and mixed with 20 µl 1,2-diamino-4,5-methylenedioxybenzene (DMB) solution, then incubated at 50°C for 2 h^32^.

For fluorometric HPLC analysis, the reaction mixtures were diluted 10-fold and directly injected onto a CAPCELL PAK C_18_ column (250 mm length × 4.6 mm i.d., Osaka Soda, Japan). Separation was performed using a mobile phase of methanol/acetonitrile/water (7:7:86, v/v/v) at a flow rate of 1 ml/min, with the column maintained at 26°C. Analysis was carried out on a Jasco LC-900 HPLC system equipped with a Jasco FP-920 fluorescence detector (excitation 373 nm; emission 448 nm). Neu5Ac, Neu5Gc (Nacalai Tesque, Japan), Kdn, and bovine submaxillary gland mucin (BP Biomedicals, LLC) were used as authentic standards.

### Lectin microarray

Lectins (see Supplementary Table S4 for list of 45 lectins) were printed in triplicates on LecChip-uni45 (Precision System Science, Japan) and activated using the manufacturer’s probing buffer. Tissue lysates from five female and five male mice (all 1 µg/µl protein concentration) were pooled prior to analysis. For quality control, 100 ng of each pooled protein sample was separated by SDS–PAGE and visualized by silver staining to confirm protein integrity and consistent loading. One microgram of protein from each pooled tissue sample was labeled with Cy3 Mono-Reactive dye (Cytiva), and the reaction quenched with Tris-buffered saline (pH 7.4). Unreacted reagents were removed using a Zeba spin desalting column (Thermo Fisher Scientific). The labeled samples were diluted with probing buffer, applied to the lectin microarray (LMA), and incubated for 16 h at 20°C. Slides were scanned using an evanescent-field excitation fluorescence imager (GlycoStation Reader 2300; GlycoTechnica) under the following conditions: exposure times of 0.5, 1, 2, 3, 4, and 5 s with a digital binning mode of 1 × 1^57^. This serial scanning generated six images per chip. The net intensity of each spot on the lectin array chip was calculated using GlycoStation Tool Pro software (v3.0).

### Transcriptomics analysis

In preparation for transcriptomics, dissected tissues were homogenized in TRI Reagent (Molecular Research Center, Inc.), immediately frozen in liquid nitrogen and stored at -80°C. Following thawing at ambient temperature, chloroform was added and centrifuged. 2-propanol was added to the aqueous phase, and RNA was extracted and quantified using the QIAGEN RNeasy Mini Kit and QunitiFluor RNA System (Promega) in accordance with the protocols provided by the manufacturer.

RNA expression was performed by low-cost and easy RNA-Seq (Lasy-Seq) and quantified, according to previous methods ^58,59^. The data was analyzed using R (v4.5.1) and edgeR (v4.6.3). Samples with a library size below 5.0×10^5^ were omitted due to low quality. Counts per million (CPM) were calculated using library sizes normalized by the trimmed mean of M-values (TMM) method implemented in edgeR.

Differential expression analysis between male and female samples was performed separately for each organ using the edgeR. Gene-wise negative binomial generalized log-linear models were fitted, and differential expression was assessed using empirical Bayes quasi-likelihood F-tests (glmQLFTest). Adjusted p value (FDR) was performed but significance was defined as *p* < 0.05. Log fold change (FC), *p*-values, and FDR were calculated using all genes as background, after which only genes annotated as glycosylation enzymes or transporters were retained and results from all organs were combined into a final table (Supplementary Table S2).

### Bioinformatic analysis

Biological process enrichment analysis based on Gene Ontology (GO) terms was performed using the Database for Annotation, Visualization, and Integrated Discovery (DAVID; National Institutes of Health), with adjusted *p* values less than 0.05 considered to be statistically significant. Subcellular localization analysis was conducted using the SubcellulaRVis online tool^60^. Differential proteome and protein-specific glycoform analyses were performed using Perseus v2.0.7.0 (https://maxquant.net/perseus/). Statistical significance was determined by t-test followed by Benjamini–Hochberg correction for multiple testing, with a false discovery rate (FDR) threshold of 0.05. Boxplots were generated using GraphPad Prism v9.4.1 (Dotmatics). If not mentioned otherwise, data are plotted as mean (average) and error bars indicate SD. Replicates and statistical significance have been stated in the figure legends.

Reduced two-dimensional principal component analysis (PCA) plots were generated for the multi-omics datasets across tissues and individual mice using the <u>pcaMethods</u> package in R. Correlation analyses were performed to assess the relationships between *N*-glycan features, lectin reactivity, and transcriptomic data across tissues. Pearson correlation coefficients were calculated, and statistically significant correlations (*p* < 0.05) were selected for visualization. The results were displayed as dot plots, where color indicates the direction (positive or negative) and size indicates the magnitude of the correlation. All correlation plots were generated using the <u>ggplot2</u> R package.

## Acknowledgments

This work was supported by KAKENHI 24K17793 (to RK), 24K08829 (to MH), 25K02224 (to CS) J-Glyconet (JGN), and Human Glycome Atlas Project (HGA). We also acknowledge the NOVARTIS foundation (Japan) for the promotion of science. KK (Kenji Kadomatsu) is supported by Japan Agency for Medical Research and Development (grant number JP25gm1410011h0004). MTA is the recipient of an Australian Research Council Future Fellowship (FT210100455).

## Data availability

The glycoproteomics and proteomics LC-MS/MS raw data have been deposited to the ProteomeXchange Consortium via the PRIDE^61^ partner repository with the dataset identifiers: PXD074760 (glycoproteomics) and PXD074840 (proteomics). Glycomics LC-MS/MS raw data were deposited to GlycoPOST with the identifiers: GPST000689.

## Author contributions

RK and KK^1^ (Kenji Kadomatsu), CS, MTA designed experiments. RK, MH, DW, FS, TN, TO, KMB, NB, MK conducted experiments. RK, KK^2^ (Ken Kitajima), CS, YM, MTA established laboratory methodologies. RK, MH, DW, TN, ZSB, SC, BZ, CNO, AK, CS analyzed the data. BZ, HK, YM developed online database. DK, KK^1^, KK^2^, CS provided reagents and intellectual input. RK, CS and MTA wrote the manuscript. RK, CS and MTA supervised this study and acquired funding. All authors have reviewed and approved the manuscript.

## Supplemental data

This article contains supplemental information including Supplementary Table S1-S9 and Supplementary Figure S1-S5.

