## Supplementary Figures for "Multi-omics definition of the sex-specific glycoproteome of murine tissues"

Running title: The sex-specific glycoproteome atlas of murine tissues

Keywords: Atlas, Glyco-enzymes, Glycoproteome, Glycosylation, Multi-omics, Sex, Tissue

***Co-corresponding authors**:

Associate Professor Rebeca Kawahara, PhD

Emergent/Innovative Engineering Building Room 815

Nagoya University, Furo-cho, Chikusa-ku, Nagoya, Aichi 464-8601, Japan

Professor Chihiro Sato, PhD

Office 520, Building B, Graduate School of Bioagricultural Science/ Institute for Glyco-core Research, Emergent/Innovative Engineering Building

Nagoya University, Chikusa, Nagoya 464-8601

Professor Morten Thaysen-Andersen, PhD

Office 333, Building 4WW, School of Natural Sciences

Macquarie University, NSW-2109, Macquarie Park - Sydney, Australia

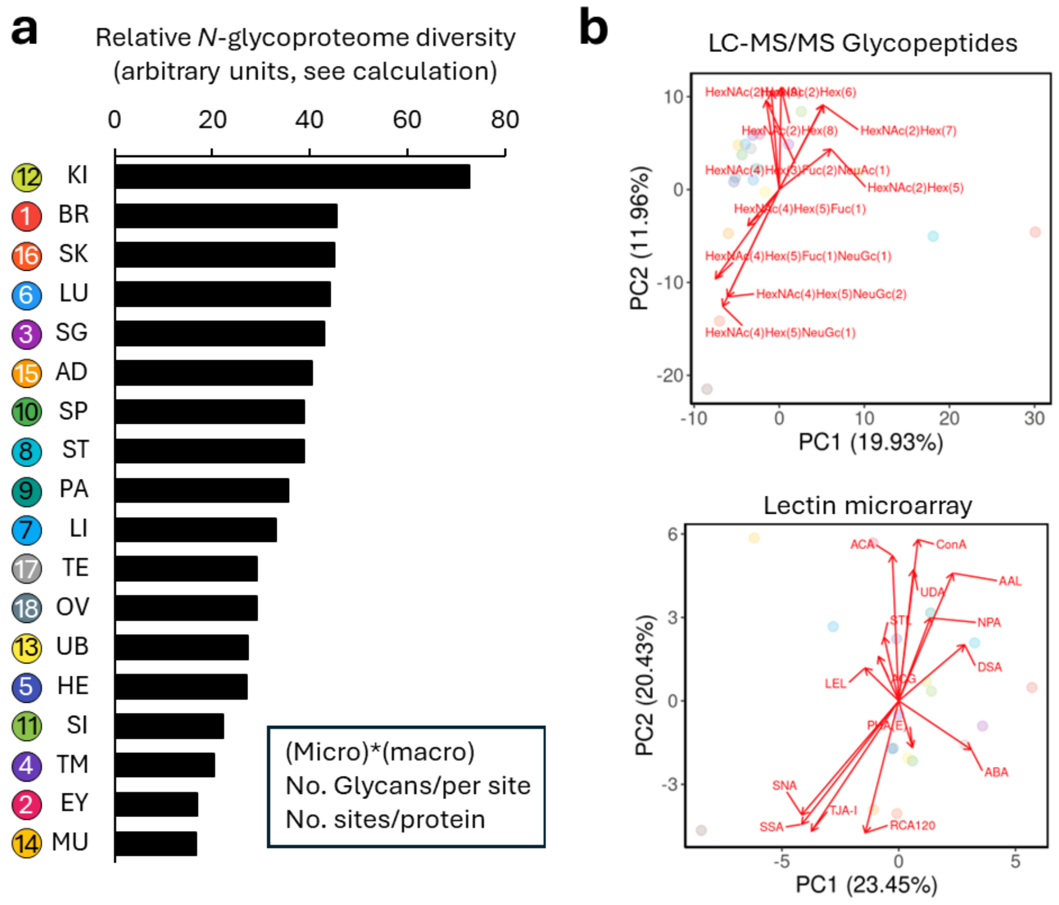


**Supplementary Figure S1. Tissue-specific *N*-glycoproteome diversity. a)** Quantification of glycoproteome diversity across tissues, calculated as the product of the number of distinct glycans identified per glycosylation site (micro-heterogeneity) and the number of occupied *N*-glycosylation sites per protein (macro-heterogeneity), expressed in arbitrary units. **b)** Multivariate biplot illustrating the contribution of glycan structural features as measured by LC-MS/MS-based glycoproteomics (top) and lectin microarray (bottom) to tissue discrimination.

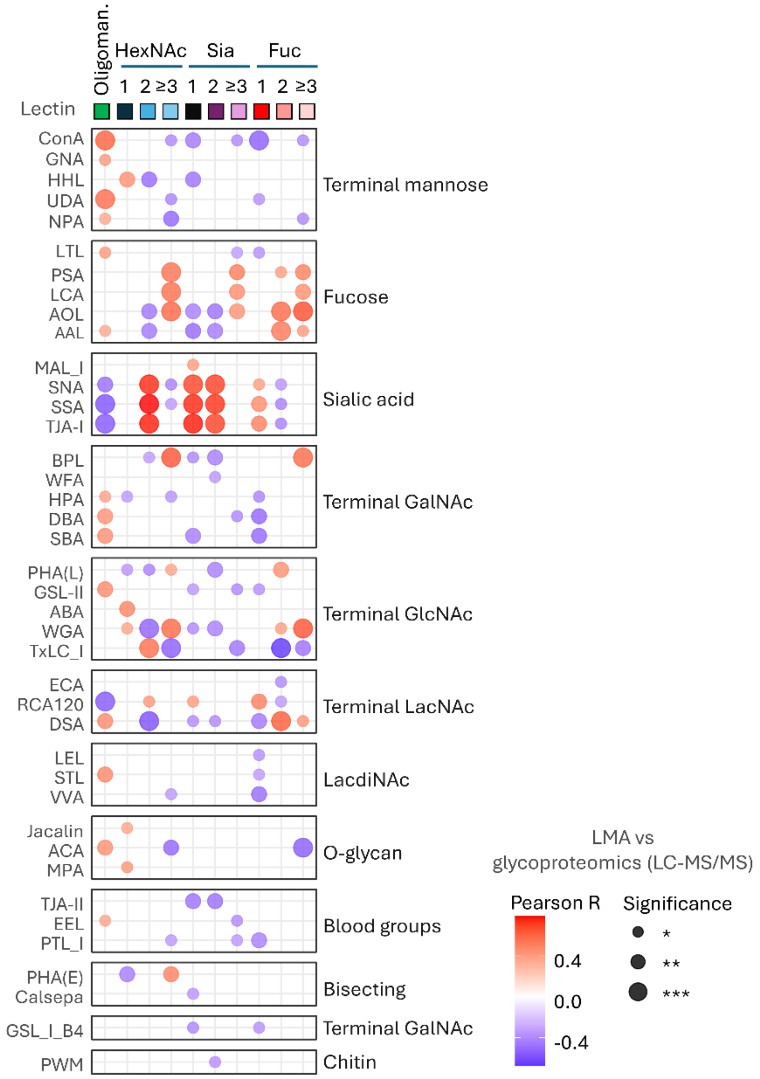


**Supplementary Figure S2. Concordance between global lectin reactivity and *N*-glycoproteomics data.** Lectins from the lectin microarray (LMA) have been grouped according to their glycan binding characteristics and correlations were investigated between their reactivity and key glycoproteome features as measured by LC-MS/MS glycoproteomics i.e. oligomannose processing, antennary branching, fucosylation and sialylation. Only significant correlations are shown (Pearson R; *$p < 0.05$, **$p < 0.01$, ***$p < 0.001$, see key for details).

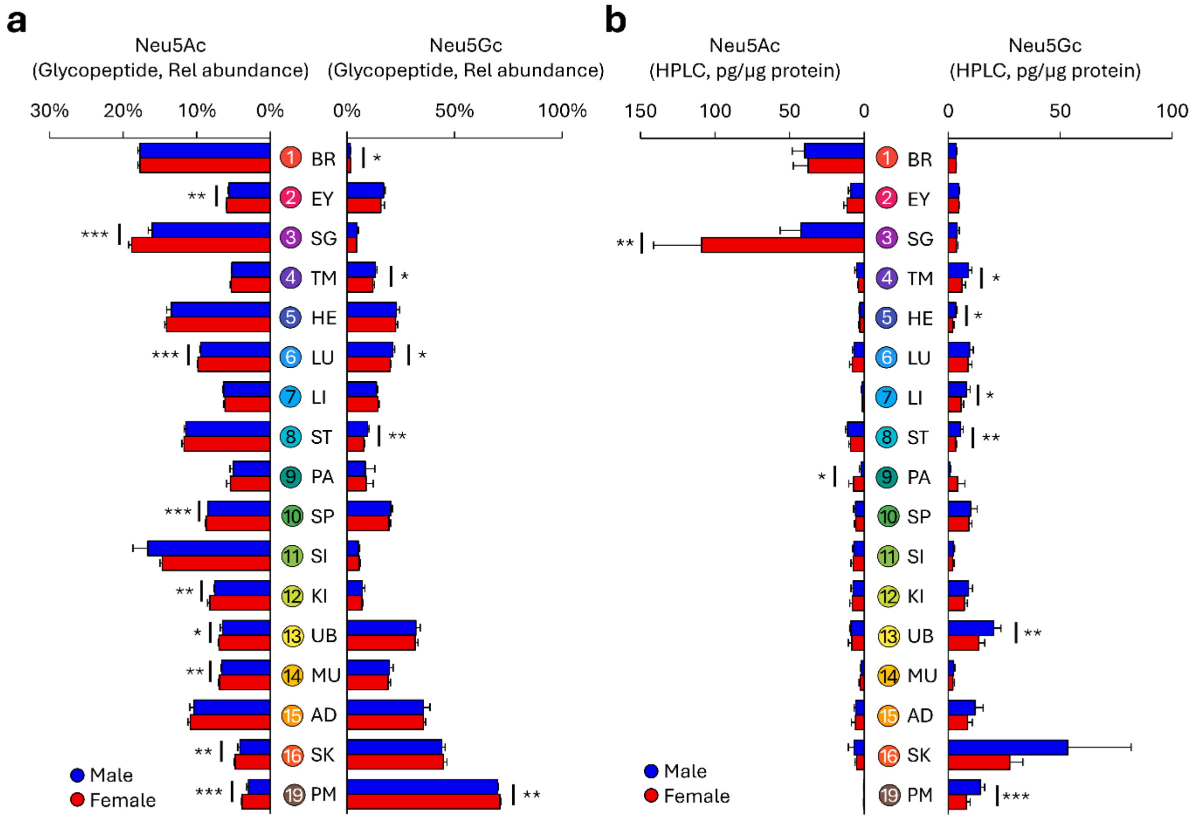


**Supplementary Figure S3. Sex-specific Neu5Ac and Neu5Gc levels across tissues. a**) LC-MS/MS-based glycopeptide quantification and **b**) HPLC-based quantification of sialic acid species across male and female tissues (n = 5 per sex) focusing exclusively on Neu5Ac and Neu5Gc. Statistical significance was determined by two-sided t-test: *$p < 0.05$, **$p < 0.01$, ***$p < 0.001$.

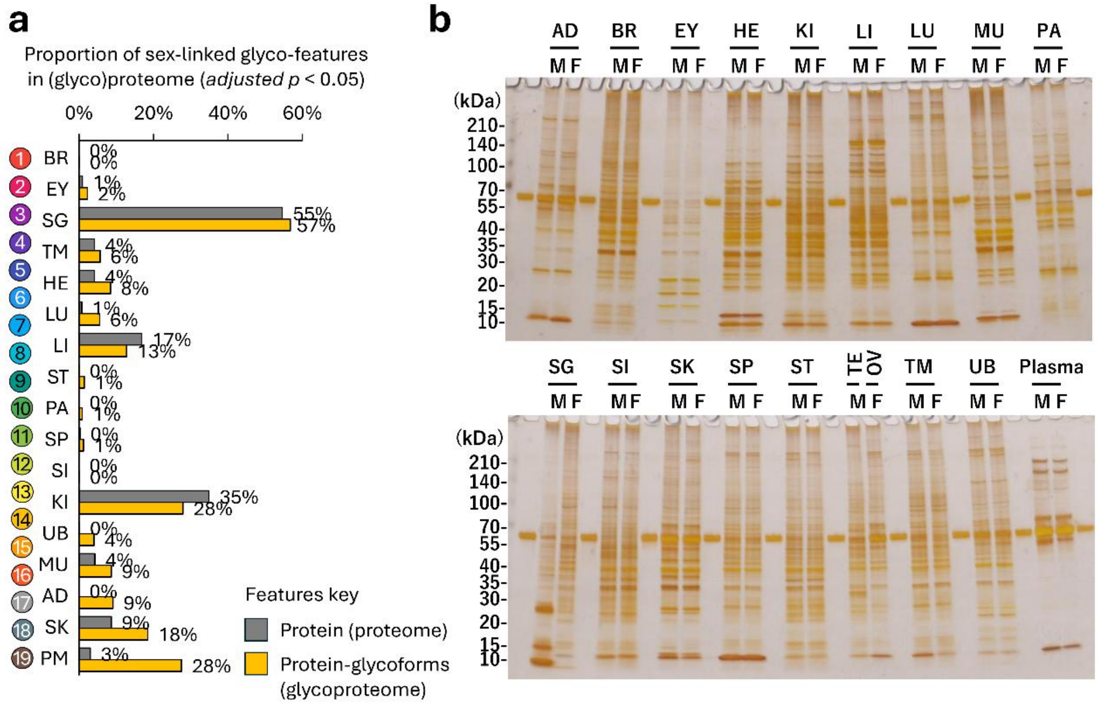


**Supplementary Figure S4. (Glyco)proteome differences across male and female mouse tissues. a)** Proportion of proteome (grey) and protein-specific glycoform (yellow) features that exhibit sex differences relative to all identifications for each tissue. Statistical analysis was performed using a two-sided t-test (n = 5 per sex) with multiple testing correction (adjusted $p < 0.05$) (see **Supplementary Table S7** and **Supplementary Table S8** for data). **b)** Representative silver-stained SDS–PAGE profiles of pooled male and female tissue lysates, with 100 ng of denatured protein loaded per lane, providing a gel-based overview of global proteome abundance patterns for each tissue.

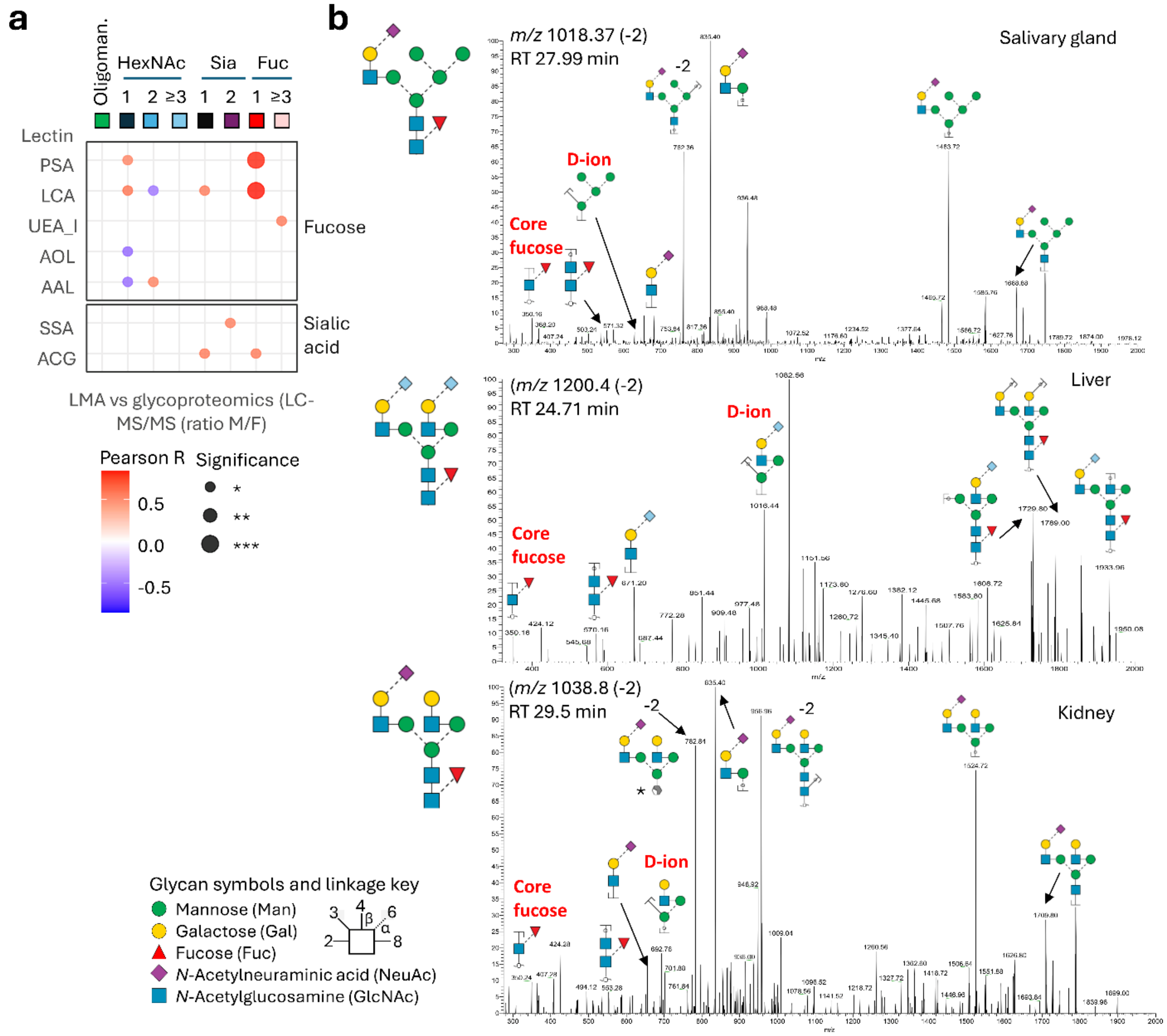


**Supplementary Figure S5. Linkage-specific analysis of fucosylation and sialylation by LMA and PGC-LC-MS/MS. a)** Correlation analysis of sex differences in glycosylation comparing global lectin binding characteristics (combined across tissues) with *N*-glycoproteome features (glycoproteomics) in males and females. Only statistically significant correlations for fucose- and sialic acid–reactive lectins are shown, see key for details. **b)** Representative PGC-LC-MS/MS spectra validating linkage-specific structural assignments, confirming the presence of α1,6-fucosylation (core fucose) and α2,6-sialylation as prominent features on *N*-glycans from the salivary gland (top), liver (middle), and kidney (bottom).
